## Supplemental Table 1 for "Comparative Analysis and Rational Design of dCas9-VP64 Variants for CRISPR Activation"

Supplemental Table 1. Key resources (Plasmids, gRNAs and primers)

| **Plasmids** | | | | | |
| --- | --- | --- | --- | --- | --- |
| **Name** | | | **Source** | | **identifier** |
| lenti dCAS-VP64_Blast | | | addgene | | # 61425 |
| Lenti-EF1α-VP64-dCas9-VP64_Blast | | | This study | | N/A |
| lentiMPH v2 | | | addgene | | # 89308 |
| lenti sgRNA(MS2)_puro backbone | | | addgene | | # 73795 |
| Lenti_MCP-VP64_Hygro | | | addgene | | # 138458 |
| dSV40-NLS-dCas9-HA-NLS-NLS-10xGCN4 | | | addgene | | # 107310 |
| EF1α-dCas9-10xGCN4_Hygro | | | This study | | N/A |
| EF1α-scFv-p65-HSF1-T2A-EGFP-WPRE-PolyA | | | addgene | | # 107311 |
| EF1α-scFv-p65-HSF1-Blast | | | This study | | N/A |
| pLV-dCas9-p300-P2A-PuroR | | | addgene | | # 83889 |
| Lenti-EF1α-dCas9-p300_Blast | | | This study | | N/A |
| Lenti-EF1α-VP64-dCas9-p300_Blast | | | This study | | N/A |
| pHRdSV40_scFv_GCN4_sfGFP_p65-hsf1_GB1_NLS | | | addgene | | # 79372 |
| Lenti-EF1α-scFv-VP64_Blast | | | This study | | N/A |
| reporter-gT1 | | | addgene | | # 47320 |
| pCR Blunt II-TOPO | | | Invitrogen | | # K275040 |
| pTRE3G | | | Takara | | # 631173 |
| pNLF1-C [CMV/Hygro] | | | Promega | | # N1361 |
| pCR-U6-gRNA-miniCMV-TdTomato | | | This study | | N/A |
| pCR-U6-gRNA-miniCMV-Nluc | | | This study | | N/A |
| pCR-U6-gRNA-TRE3G-TdTomato | | | This study | | N/A |
| pCR-U6-gRNA-TRE3G-Nluc | | | This study | | N/A |
| pGL4.54 [luc2/TK] | | | Promega | | E5061 |
| **gRNA** | | | | | |
| **Gene** | | **Sequence** | **Reference** | | |
| hASCL1 | | GCAGCCGCTCGCTGCAGCAG | Nature 517,7536, 583-8 (2015) | | |
|  | | TGGAGAGTTTGCAAGGAGC | Nat Methods 10, 973–976 (2013) | | |
|  | | GTTTATTCAGCCGGGAGTC | Nat Methods 10, 973–976 (2013) | | |
| hMYOD1 | | GGGCCCCTGCGGCCACCCC | Nature 517,7536, 583-8 (2015) | | |
|  | | CTCCCTCCCTGCCCGGTAG | Nat Biotechnol 33, 510–517 (2015) | | |
|  | | AGGTTTGGAAAGGGCGTGC | Nat Biotechnol 33, 510–517 (2015) | | |
| hNEUROD1 | | GAGGTCCGCGGAGTCTCTAAC | Nat Methods 13, 563–567 (2016) | | |
| hIL1RN | | GCATCAAGTCAGCCATCAGC | Nat Biotechnol 33, 510–517 (2015) | | |
|  | | TGTACTCTCTGAGGTGCTC | Nat Biotechnol 33, 510–517 (2015) | | |
|  | | ACGCAGATAAGAACCAGTT | Nat Biotechnol 33, 510–517 (2015) | | |
| mNeurog2 | | TGGTTCAGTGGCTGCGTGTC | Nature 517,7536, 583-8 (2015) | | |
| mHbb-bh1 | | AGAGAGTCTGGGCAAGACAG | Nat Methods 13, 563–567 (2016) | | |
| miniCMV | | GTCCCCTCCACCCCACAGTG | Nat Methods 12, 326-8 (2015) | | |
| TRE3G | | TACGTTCTCTATCACTGATA | Nature methods 13,1043-1049 (2016) | | |
| **Primers** | | | | | |
| **Gene** | **Sequence (5’-3’)** | | **Strand** | **Reference** | |
| hASCL1 | CGGTCTCATCCTACTCGTCG | | sense | This study | |
|  | GTTGTGCGATCACCCTGCT | | antisense | This study | |
| hMYOD1 | TCCGACGGCATGATGGACTA | | sense | This study | |
|  | CAGTCTAGGCTCGACACCG | | antisense | This study | |
| hNEUROD1 | CAGGACCTACTAACAACAAAGGAAA | | sense | This study | |
|  | GACACTCGTCTGTCCAGCTT | | antisense | This study | |
| hIL1RN | GGAATCCATGGAGGGAAGAT | | sense | Nat Biotechnol 33, 510–517 (2015) | |
|  | TGTTCTCGCTCAGGTCAGTG | | antisense | Nat Biotechnol 33, 510–517 (2015) | |
| hACTB | TGGCACCCAGCACAATGAA | | sense | JASN 30, 304-321 (2019) | |
|  | CTAAGTCATAGTCCGCCTAGAAGCA | | antisense | JASN 30, 304-321 (2019) | |
| mNeurog2 | CAAAGTCGCCCAGCCGAGA | | sense | This study | |
|  | GATTTGACGAACATCCTACGC | | antisense | This study | |
| mHbb-bh1 | CTGGGAAGGCTCCTGATTGT | | sense | Nat Methods 13, 563–567 (2016) | |
|  | GTTCTTAACCCCCAAGCCCA | | antisense | Nat Methods 13, 563–567 (2016) | |
| mGapdh | CCTGGAGAAACCTGCCAAGTATG | | sense | This study | |
|  | GGTCCTCAGTGTAGCCCAAGATG | | antisense | This study | |
